# Closed Kinematic Chain Biomechanics and Cycling: Linking Biomechanical Variables to Knee Joint Loading

**DOI:** 10.64898/2026.03.29.715123

**Authors:** Honore Baho Vita, Dawit Fsaha Welegebriel

## Abstract

This study investigates closed kinematic chain biomechanics in cycling with a focus on knee joint loading. Data from 16 cyclists collected on a standardized ergometer were analyzed in OpenSim using inverse dynamics, static optimization, and joint reaction analysis. To keep the pipeline consistent across all subjects, the report summarizes right-knee outputs over a steady-state interval between 120 and 124 s. Peak knee joint moments ranged from 15.79 to 44.85 Nm (mean 30.49 ± 7.66 Nm), while peak resultant knee reaction forces ranged from 1187.61 to 3309.04 N (mean 2317.19 ± 728.19 N). Static optimization showed strong contributions from the rectus femoris and vastus lateralis during power production, with additional stabilization from the biceps femoris long head and gastrocnemius medialis. Mean peak muscle activation was highest for the rectus femoris (0.72 ± 0.19), followed by the biceps femoris long head (0.66 ± 0.20). Mean peak muscle force was highest for the vastus lateralis (1078.1 ± 305.8 N) and rectus femoris (994.1 ± 379.2 N). The results confirm substantial inter-subject variability in knee loading and support the use of personalized training or rehabilitation strategies when cycling is used for performance development or joint recovery.

## I. INTRODUCTION

Cycling is recommended for the early stages of post-operative or post-traumatic rehabilitation of the musculoskeletal system since it exposes the injured joint to a relatively low workload [1]. In research, it is used to better understand the mechanical aspects of human movements where it provides unique biomechanical challenges and opportunities for examining human motion dynamics. Numerous studies focus on Closed Kinematic chains (CKCs), in which joint movements are interconnected such that movement at one joint directly influences others [2]. Therefore, CKC analysis helps determine the forces on joints at different angles, assess muscle activation patterns, and prevent movements that may lead to injuries [3]. Advancements in technology, especially in computational modeling, have transformed the investigation of CKC systems, with computer simulations widely used to run tests and reduce the time and cost of developing new mechanical systems. This paper aims to explore the biomechanical principles of CKCs in cycling by leveraging simulation tools such as OpenSim and MoCo. We focus on analyzing the biomechanical interactions between bicycle and rider during cycling, with the goals of enhancing cyclist performance, accelerating rehabilitation, and minimizing injury risks. This research integrates data from 16 cyclists of varying experience levels, captured using motion capture technology and force sensors, providing valuable insights that can lead to personalized training programs and interventions designed to improve cycling efficiency and safety.

## II. RELATED WORK

Researchers use mathematical models in cycling biomechanics to investigate the forces and torques in the lower extremities during pedaling. Notably, several studies have used inverse dynamics simulations to assess joint contact forces and net moments, providing insights into potential injury processes and prevention [4]. Similarly, Handke and Balchanowski used inverse kinematics simulation to create movement parameters for all lower limb joints [5]. According to their results, they were able to simulate active moments for the hip joints in the sagittal plane which they claim was not possible to simulate an individual joint. Extending their findings, Handke and Balchanowski (2019) carried out research on the constrain between the foot and pedal when cycling [6]. According to their study, there was no significant difference in the efficacy of torque generated when using cleats to fix the foot and not affixing the foot to the pedal. Thus, they focused on the relationship between pedal and foot to determine the displacement characteristics in specific lower limb joints at a given cadence.

Furthermore, biomechanical models are used to estimate the joint forces and according to Bini et al. (2014), this is difficult because muscle forces change with muscle length, joint angles change throughout the pedaling cycle, and the locations of the crank relative to leg segments determine the leverage of each muscle, the forces applied to the pedals fluctuate, resulting in a mismatch between force direction and effective crank propulsion [4]. The authors stress that biomechanical and mechanical modeling approaches are used to quantify forces operating on body segments to prevent cycling accidents. Cycling efficiency is affected by various biomechanical parameters such as the pedaling frequency, the saddle height, the crank-arm length, and the seat tube angle [7]. Therefore, for a biomechanical analysis of cycling both the theoretical model and a well-equipped device are required to obtain the parameters necessary to monitor the activity. Thus, the movement and forces involved in the lower limbs and bike crank can be examined using the mathematical model. Generally in engineering, computer simulations are widely used to run tests and reduce the cost and time of new mechanical systems developments. Newton-Euler equations of a rigid body are one of the approaches mostly used to represent multibody systems mathematically [8].

A kinematic chain is a result of linking rigid bodies through mechanical constraints [9]. Escamilla et al. (1998) cite research indicating that physical exercises can be classified into two categories; Closed Kinematic Chain (CKC) and Open Kinematic Chain (OKC) exercises [10]. As stated in their study, OKC exercises are when the distal part of the body is free of movement without resistance whereas the exercises are considered to be CKC if there is an external movement resistance. The authors emphasize that CKC exercises are often recommended to people after reconstruction surgery of the anterior cruciate ligament (ACL). Patients who are in the early phases of rehabilitation benefit from CKC activities like cycling since it’s important to maintain the affected joints under minimal strain [1]. Closed kinetic Chain analysis is important in determining forces that are acting on the joints at different angles, assessing muscle activation patterns, and determining and preventing movements that may lead to injuries [3].

Zangerl and Steinicke (2021) examine the structure 3D CKC system by analyzing the joint angles of its spherical joints [11]. They stress that it is crucial to understand the configuration spaces of CKCs since they generally appear in various scientific domains including computer graphics, robotics, physics, computational biology, and protein kinematics. Similarly, Olsen (2019) asserts that CKCs are found throughout musculoskeletal systems [12]. He clarifies that the CKCs in musculoskeletal systems can be momentarily as an individual comes in contact with something or the permanent structures consist of skeletal elements, muscles, and ligaments. He states that CKCs have been the main model utilized in previous research to study the transmission of force and motion in musculoskeletal systems.

According to this study, CKCs have limited degrees of freedom (DoF) compared to OKCs and the author argues that the effect of CKCs in reducing freedom of movement has been disregarded and unexplored in a variety of musculoskeletal systems where many studies focus on permanent CKCs structures. Based on two characteristics of the linkage mobility formula the number of loops and the type of constituent joint mobilities Olsen (2019) proposed classifying the mobility of biological CKCs. He uses terms like “constant” and “conditional” to describe parts of joint mobilities that are constant over variable, and “permanent” and “transient” to describe a constant vs varying number of loops, respectively. He adds that this classification is significant because it distinguishes the causes of dynamic variations in mobility during and between activities.

Analyzing the biomechanics of cycling workouts using a CKC system involves both elements of the rider and the bicycle [13]. Zwieten et al. (2016) claim that these elements are the frame, the crank, the pedal and foot, the lower leg, the upper leg, and the pelvis. Correspondingly, Malfait et al. (2010) propose that the bicycle-rider CKC system can be represented by a five-bar linkage model consisting of the thigh, the shank, the foot, the crank, and the linkage crank axis, and the hip joint [14]. This system consists of two fixed pivot joints (the crank axis and the hip joint) and three movable pivot points (the pedal spindle, the ankle joint, and the knee joint). When in motion, this CKC system is controlled and stabilized by three extensors and flexors antagonistic muscle groups of the hip, knee, and ankle [13]. Malfait et al. (2010) claim that only crank angle and pedal angle variables are required to define five-bar linkage as a kinematic system. In their study, they express the pedal (foot) angle as a function of the crank angle.

Many studies explored different approaches to advance our understanding of cycling dynamics, optimizing performance and injury prevention strategies. Cycling biomechanical modeling and analysis investigate the mechanical components of human movements, especially how muscle forces, joint torques, and limb kinematics contribute to efficient pedaling mechanics. In simulations such as inverse dynamics, these models help to estimate joint kinematics which can be used to identify changes in joint forces in injured cyclists [4]. However, they often rely on assumptions due to a lack of in vivo experimental data on physical properties and muscle activation patterns [15] and fail to capture the complex human movement dynamics. More research in this area could include real-time feedback systems and detailed muscle activation patterns to create models with greater accuracy. Bini et al. (2014) claim that there is limited research on 3D motion analysis and pedaling irregularities. Therefore, they propose that further evaluation of muscle activation and pedal forces for a more thorough understanding of cyclists’ coordinative patterns. Furthermore, According to the authors, there have been no performance improvements by changing the bike’s configuration or adjusting the pedal force direction. There is insufficient research linking these factors to injury occurrence, and preventive interventions are primarily based on empirical information. Thus, they imply the requirement for dynamic monitoring of joint kinetics and kinematics for injury prevention.

## III. METHODS

This study used data from a previous experiment involving 16 participants with different levels of cycling experience. Participants performed cycling trials on a standardized ergometer instrumented with motion capture markers and pedal force measurements. The available processed files for each subject were organized into three OpenSim output sets: inverse dynamics (ID), static optimization (SO), and joint reaction analysis (JRA).

Inverse dynamics was used to estimate the net joint moments required to reproduce the recorded motion. For consistency across all subjects, the present report focused on the right knee and extracted the knee_angle_r_moment output from each subject’s ID file. The force-like beta terms reported by the same ID files were not interpreted as physical knee joint reaction forces because they represent generalized outputs rather than internal joint loading.

Joint reaction analysis was then used to quantify internal knee joint loading. For each subject, the right-knee reaction force components expressed in the tibial frame (walker_knee_r_on_tibia_r_in_tibia_r_fx, fy, and fz) were combined into a resultant force magnitude using the square root of the sum of squared components. Static optimization files were used to extract muscle activation and force histories for four muscles relevant to cycling mechanics: rectus femoris, vastus lateralis, biceps femoris long head, and gastrocnemius medialis.

A steady-state interval between approximately 120 and 124 s was used for waveform visualization and subject comparison. For each subject, peak absolute knee moment and peak resultant knee reaction force were calculated. Subjects were also grouped into low, medium, and high loading bands based on peak moment or peak force so that waveform differences could be compared without plotting all 16 traces on the same axis.

Descriptive statistics were computed across subjects for the knee moment and knee reaction force summaries. For the muscle analyses, mean peak activation and mean peak force were calculated across subjects. Waveforms were visualized using mean ± 1 SD plots and grouped overlays to show both central tendency and inter-subject variability.

## IV. RESULTS

### A. INVERSE DYNAMICS RESULTS

Inverse dynamics results showed clear between-subject variability in right-knee loading. Peak absolute knee moment values ranged from 15.79 to 44.85 Nm, with a group mean of 30.49 ± 7.66 Nm. The highest peak moments were observed for P10 (44.85 Nm), P13 (39.46 Nm), and P06 (39.42 Nm). The lowest values occurred in P11 (15.79 Nm), P04 (20.63 Nm), and P14 (22.03 Nm).

The mean ± SD waveform highlights a repeating oscillatory pattern through the steady-state interval, while the grouped plots show that the high-moment subjects produce larger positive and negative excursions than the low-moment subjects. This indicates that net knee loading in cycling is not uniform across riders, even when the task and experimental setup are standardized.

**Table 1.** Peak right-knee moment and peak right-knee reaction force for each participant.

| Participant | Peak knee moment (Nm) | Peak knee reaction force (N) |
| --- | --- | --- |
| P01 | 25.11 | 1388.02 |
| P02 | 30.91 | 3159.99 |
| P03 | 26.3 | 1187.61 |
| P04 | 20.63 | 1696.4 |
| P05 | 28.85 | 3203.47 |
| P06 | 39.42 | 3141.92 |
| P07 | 32.13 | 3240.73 |
| P08 | 35.71 | 2006.14 |
| P09 | 31.54 | 1996.68 |
| P10 | 44.85 | 2366.1 |
| P11 | 15.79 | 2016.5 |
| P12 | 35.62 | 2076.62 |
| P13 | 39.46 | 3309.04 |
| P14 | 22.03 | 2008.5 |
| P15 | 26.52 | 1465.11 |
| P16 | 32.91 | 2812.2 |

**Figure 1.**
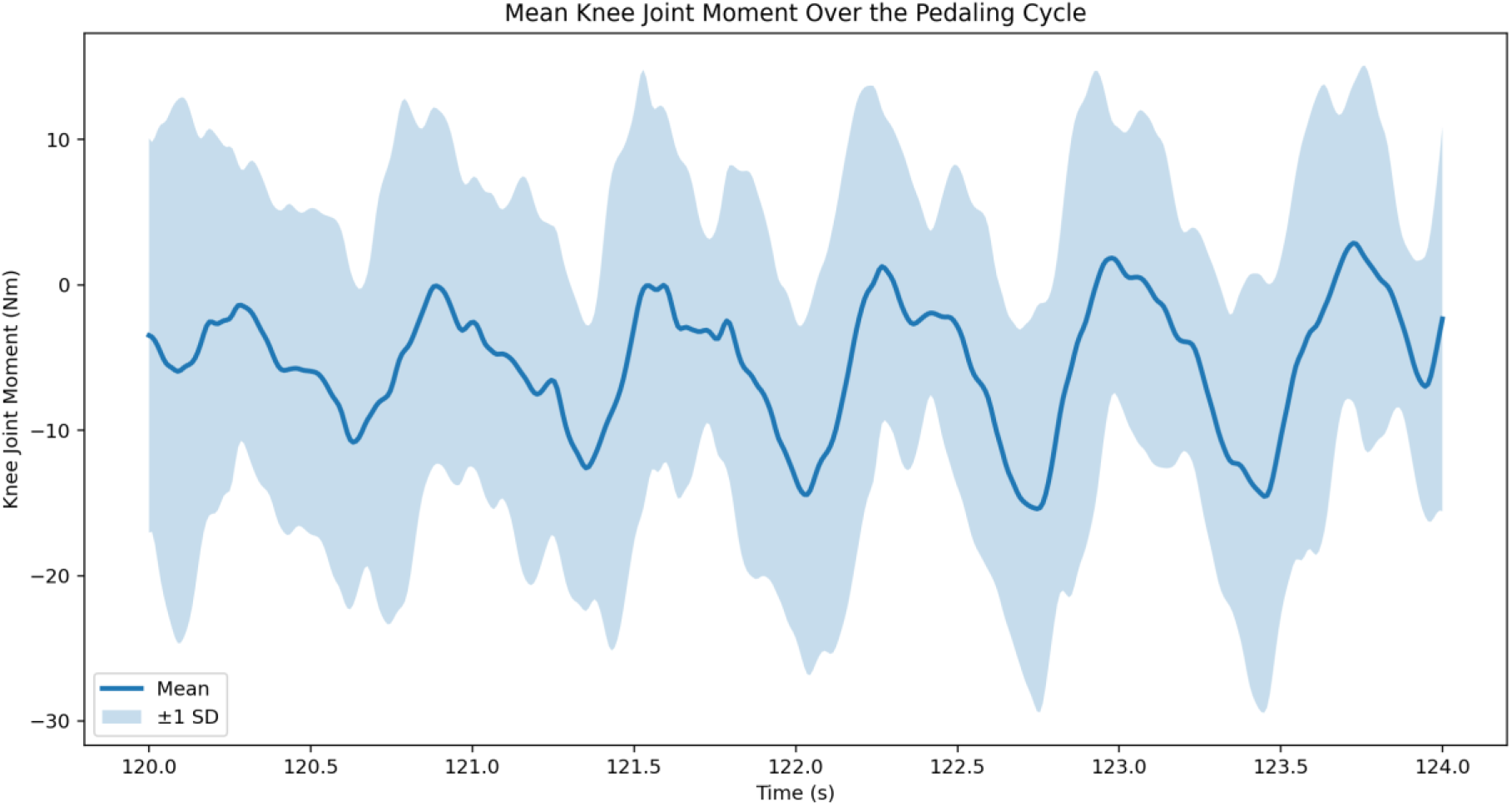
Mean ± 1 SD right-knee moment across the steady-state interval.

**Figure 2.**
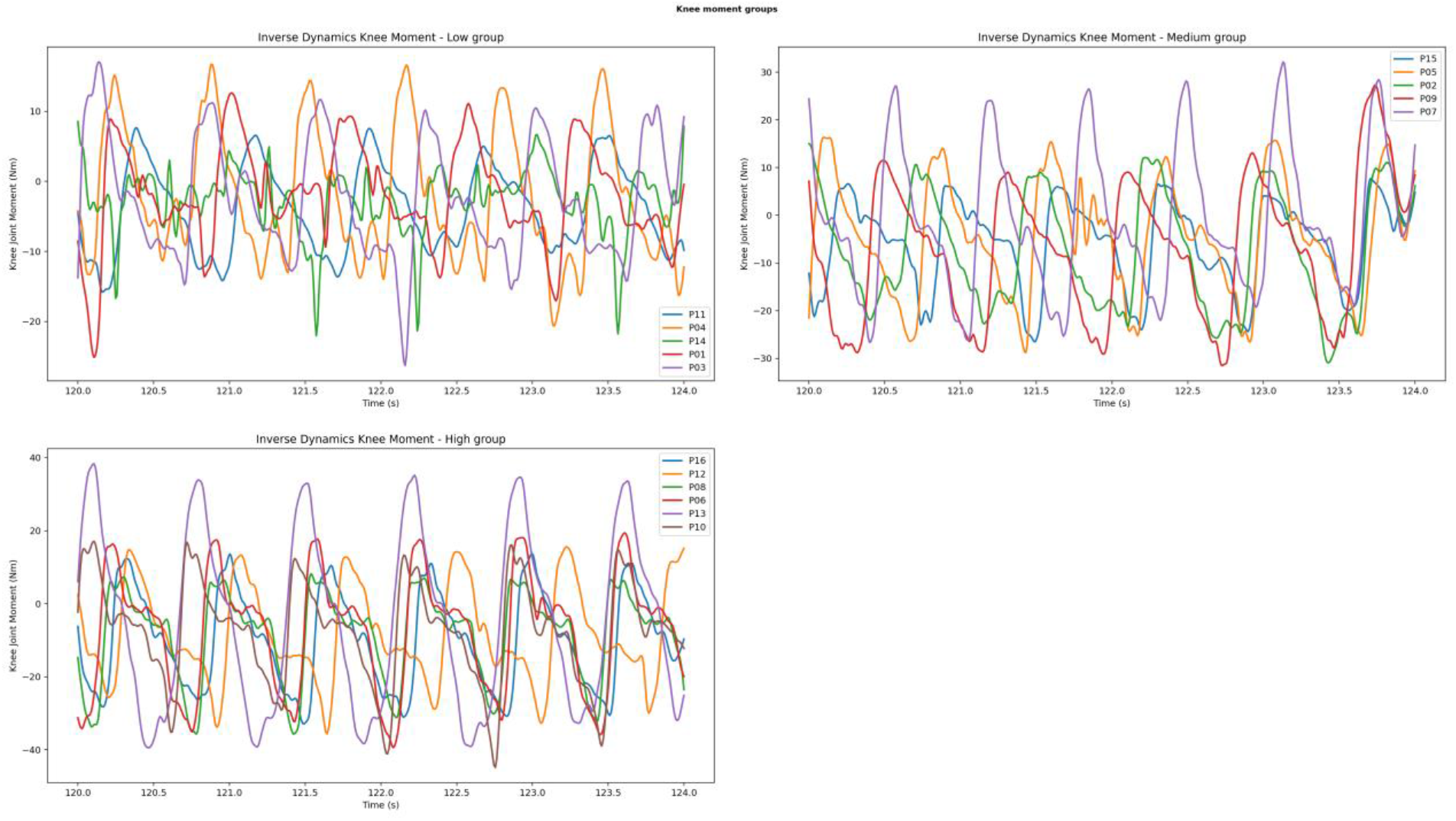
Grouped right-knee moment waveforms for low, medium, and high peak-moment subjects.

### B. JOINT REACTION ANALYSIS RESULTS

Joint reaction analysis provided the internal knee loading estimates that should be used when discussing true knee joint force. Peak resultant right-knee reaction forces ranged from 1187.61 to 3309.04 N, with a mean of 2317.19 ± 728.19 N. The highest peak reaction forces were found in P13 (3309.04 N), P07 (3240.73 N), and P05 (3203.47 N). The lowest peaks occurred in P03 (1187.61 N), P01 (1388.02 N), and P15 (1465.11 N).

Compared with the inverse-dynamics moments, the reaction force waveforms showed a wider spread across subjects and more pronounced sharp peaks. The mean ± SD curve suggests that substantial compressive and shear demands recur throughout the loaded phases of pedaling, while the grouped plots show that high-force riders maintain consistently larger force amplitudes across the full steady-state interval.

**Figure 3.**
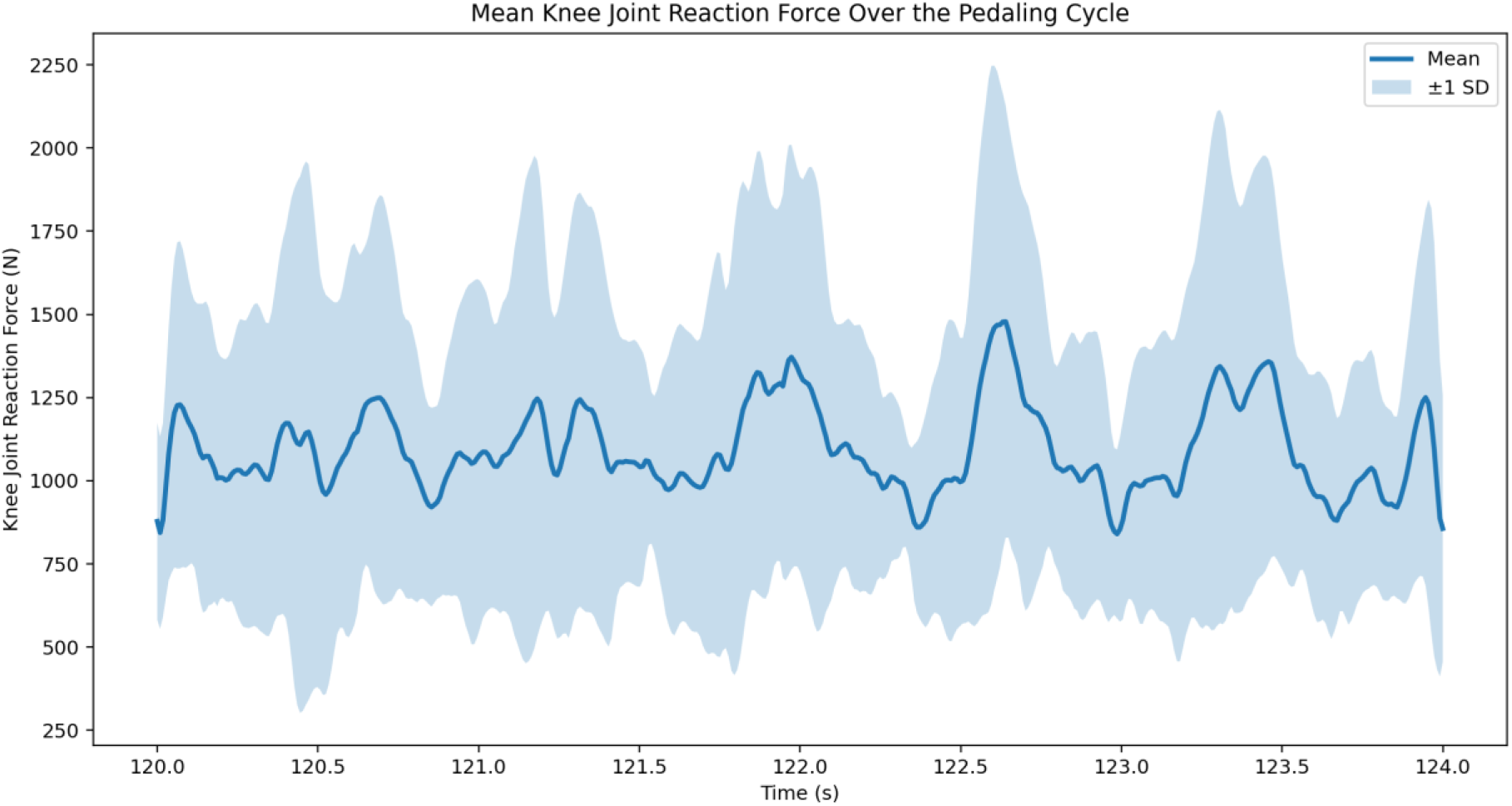
Mean ± 1 SD right-knee reaction force across the steady-state interval.

**Figure 4.**
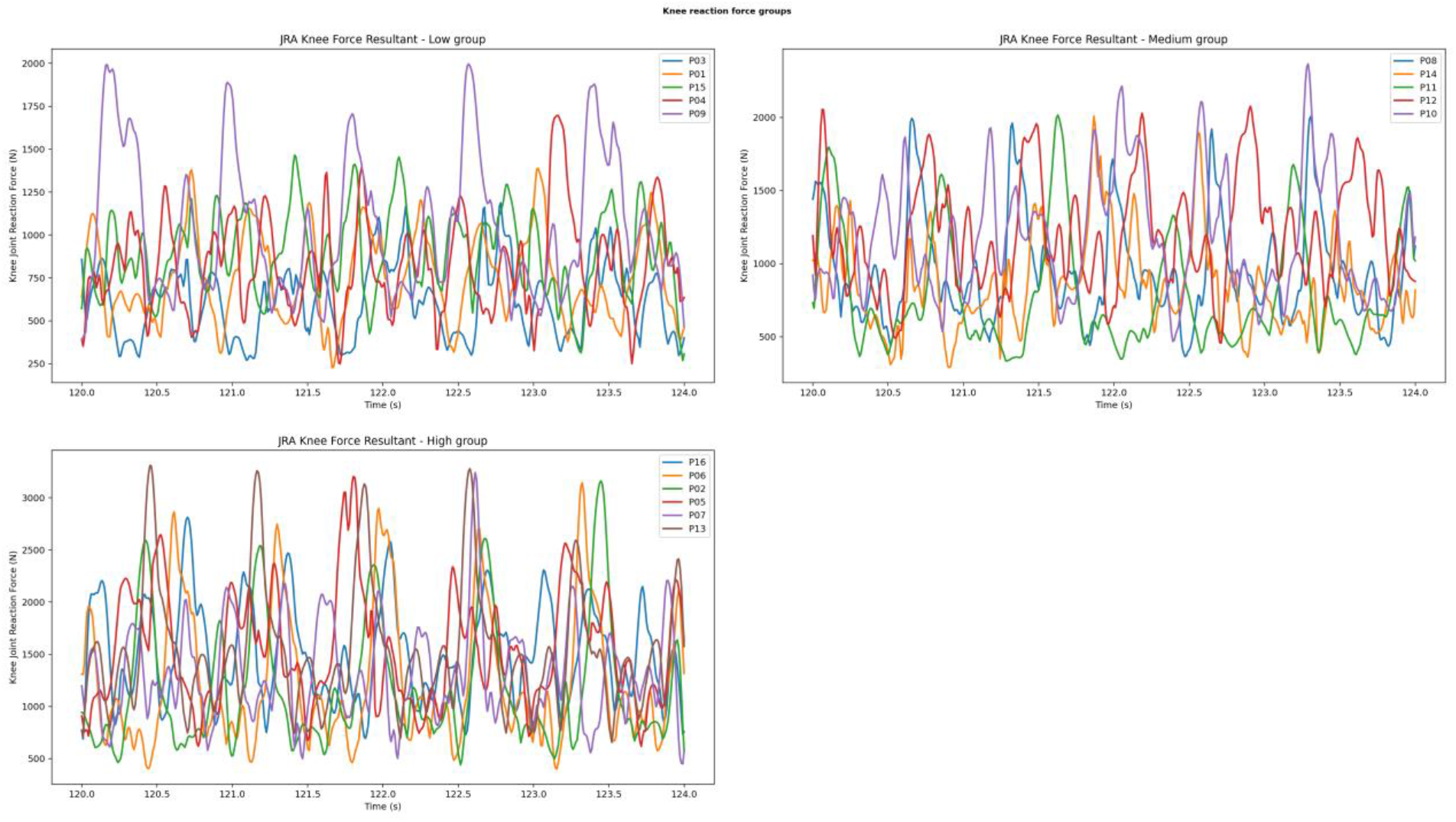
Grouped right-knee reaction force waveforms for low, medium, and high peak-force subjects.

### C. STATIC OPTIMIZATION RESULTS

Static optimization results identified a quadriceps-dominant pattern for power production, with hamstrings and gastrocnemius contributing to control and stabilization. Across subjects, mean peak activation was highest for the rectus femoris (0.72 ± 0.19), followed by the biceps femoris long head (0.66 ± 0.20), the gastrocnemius medialis (0.38 ± 0.23), and the vastus lateralis (0.34 ± 0.09).

Mean peak muscle force was highest for the vastus lateralis (1078.1 ± 305.8 N) and rectus femoris (994.1 ± 379.2 N), followed by the biceps femoris long head (771.7 ± 249.9 N) and gastrocnemius medialis (448.9 ± 192.6 N). These values support the interpretation that the quadriceps provide the main extensor demand during cycling, while the hamstrings and gastrocnemius modulate joint stability and help coordinate the transition between loaded and unloaded phases.

Across the 16 subjects, mean peak activation values were 0.72 ± 0.19 for the rectus femoris, 0.34 ± 0.09 for the vastus lateralis, 0.66 ± 0.20 for the biceps femoris long head, and 0.38 ± 0.23 for the gastrocnemius medialis. Mean peak force values were 994.1 ± 379.2 N, 1078.1 ± 305.8 N, 771.7 ± 249.9 N, and 448.9 ± 192.6 N for the same muscles, respectively.

**Figure 5.**
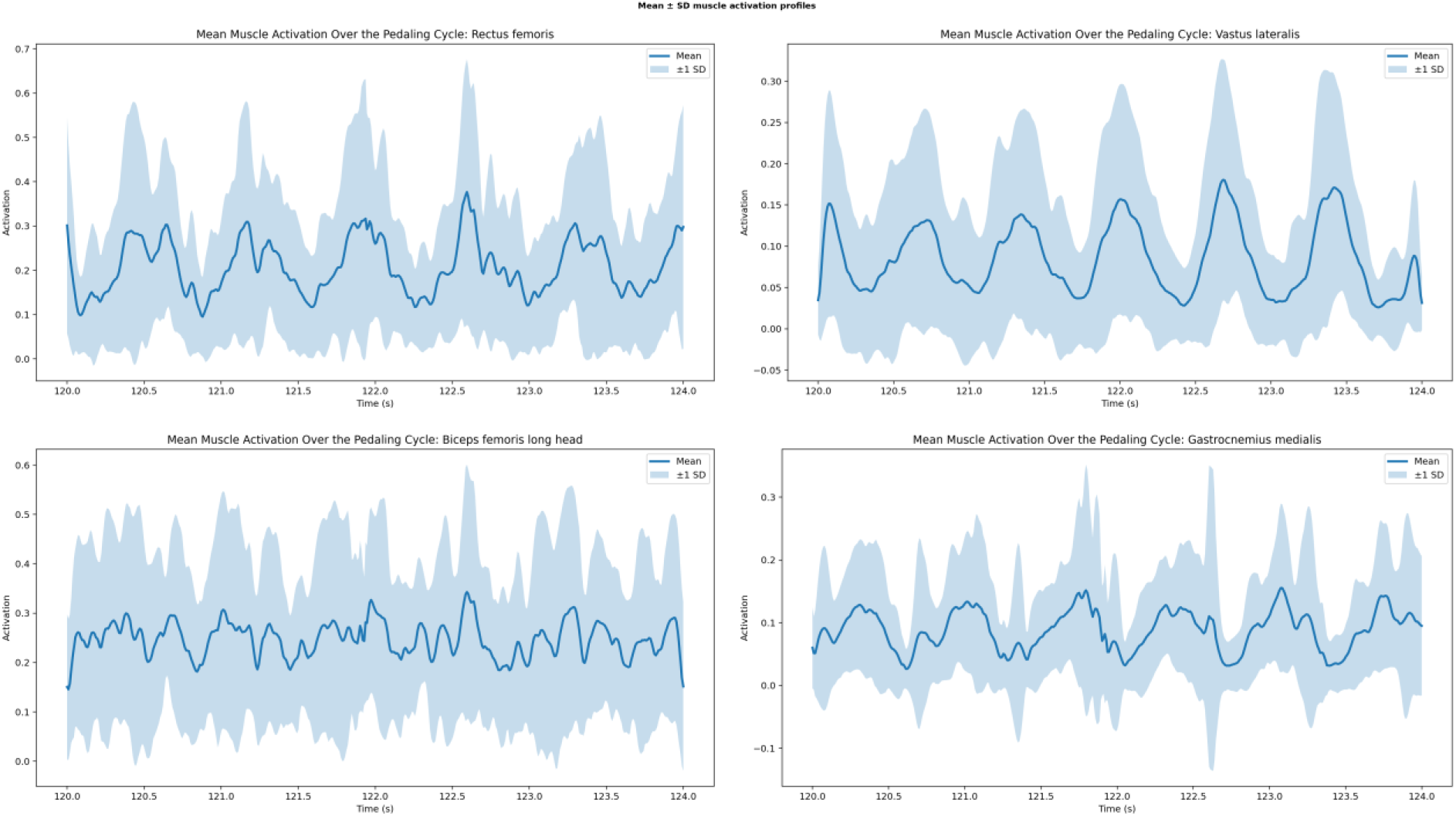
Mean ± 1 SD activation profiles for the rectus femoris, vastus lateralis, biceps femoris long head, and gastrocnemius medialis.

**Figure 6.**
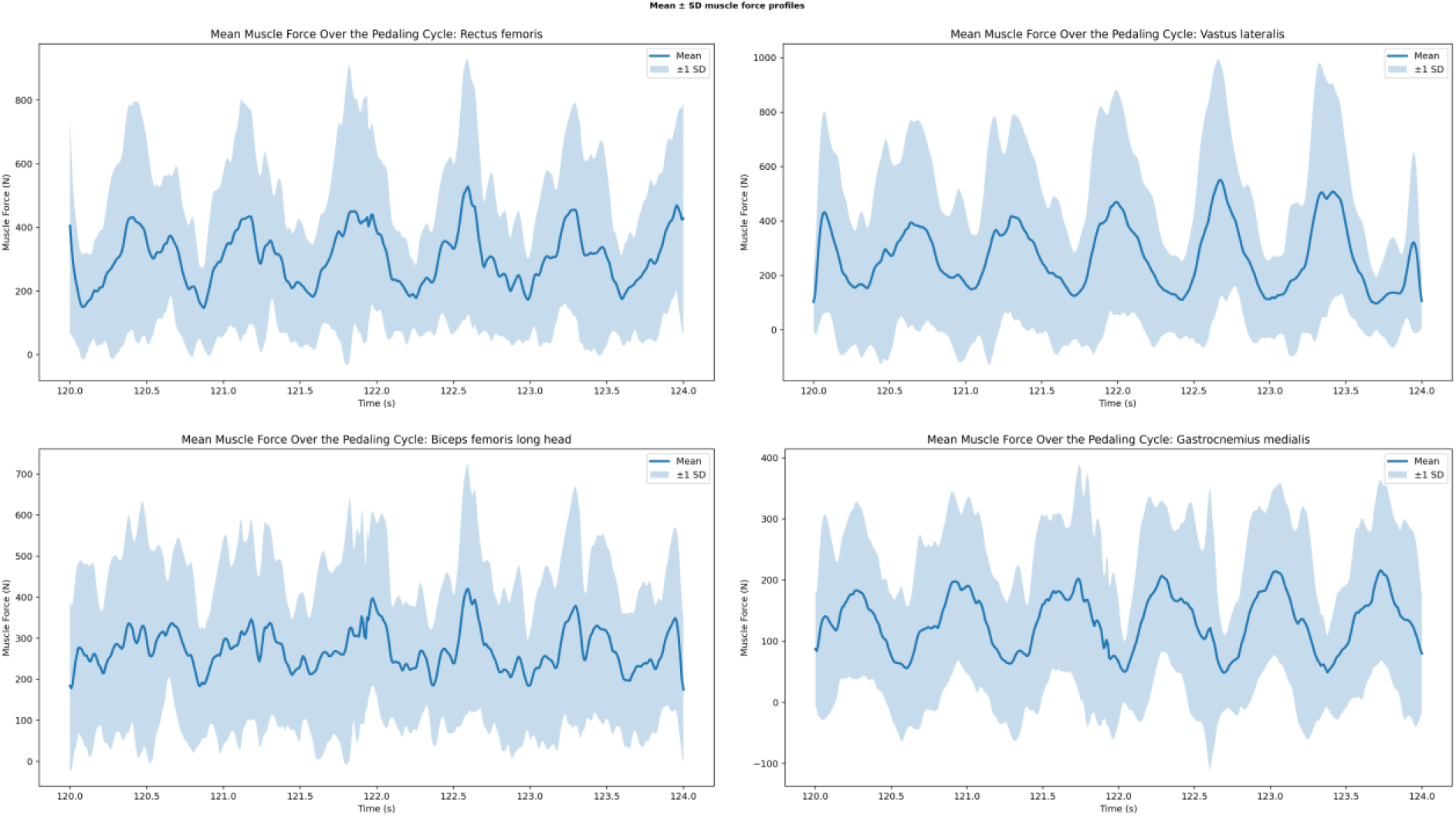
Mean ± 1 SD force profiles for the rectus femoris, vastus lateralis, biceps femoris long head, and gastrocnemius medialis.

## V. DISCUSSION

The results show a clear picture of cycling knee biomechanics under a closed kinematic chain task. Net knee moments were in the tens of Newton-meters, while internal joint reaction forces were in the low-thousands of Newtons. This distinction matters because inverse dynamics and joint reaction analysis answer different questions: inverse dynamics describes the net moment required to reproduce the motion, whereas joint reaction analysis estimates the internal force transmitted through the joint.

The results reinforce the importance of inter-subject variability. The difference between the lowest and highest peak knee moments (15.79 vs 44.85 Nm) and between the lowest and highest peak reaction forces (1187.61 vs 3309.04 N) suggests that riders performing the same task can still experience meaningfully different mechanical demands. From a practical standpoint, this supports individualized bicycle fitting, cadence control, strength work, and rehabilitation progression rather than assuming one loading profile fits all riders.

The muscle results help explain these loading differences. Rectus femoris and vastus lateralis produced the largest mean peak forces, which is consistent with their role in knee extension and power transfer during cycling. At the same time, the biceps femoris long head and gastrocnemius medialis remained active enough to support joint control, particularly when the external loading pattern changed over time. This balance between power generation and stabilization is likely one reason some riders show smoother waveforms than others.

A strength of this analysis is that it uses one consistent OpenSim pipeline across all 16 subjects and separates the role of each tool correctly. A limitation is that the present report summarizes the right knee only, and it relies on model-based estimates rather than direct in vivo measurements or EMG validation. In addition, the reported waveforms come from a selected steady-state interval rather than cycle-normalized averages, so future work should combine time-normalized ensemble analysis with 3D kinematics and subject-specific clinical or performance outcomes.

## VI. CONCLUSION

This study shows that cycling produces measurable and participant-specific changes in right-knee moment, internal reaction force, muscle activation, and muscle force. Peak knee moments ranged from 15.79 to 44.85 Nm, while peak knee reaction forces ranged from 1187.61 to 3309.04 N across the 16 cyclists. Quadriceps muscles, especially the rectus femoris and vastus lateralis, dominated power production, while the biceps femoris long head and gastrocnemius medialis contributed to control and stabilization. Together, these results support the use of personalized training and rehabilitation strategies when cycling is used either to improve performance or to manage knee joint loading during recovery.

## APPENDIX

Detailed mean ± SD activation and force profiles for the four muscles selected for the static optimization analysis are provided below.

**Appendix Figure A1.**
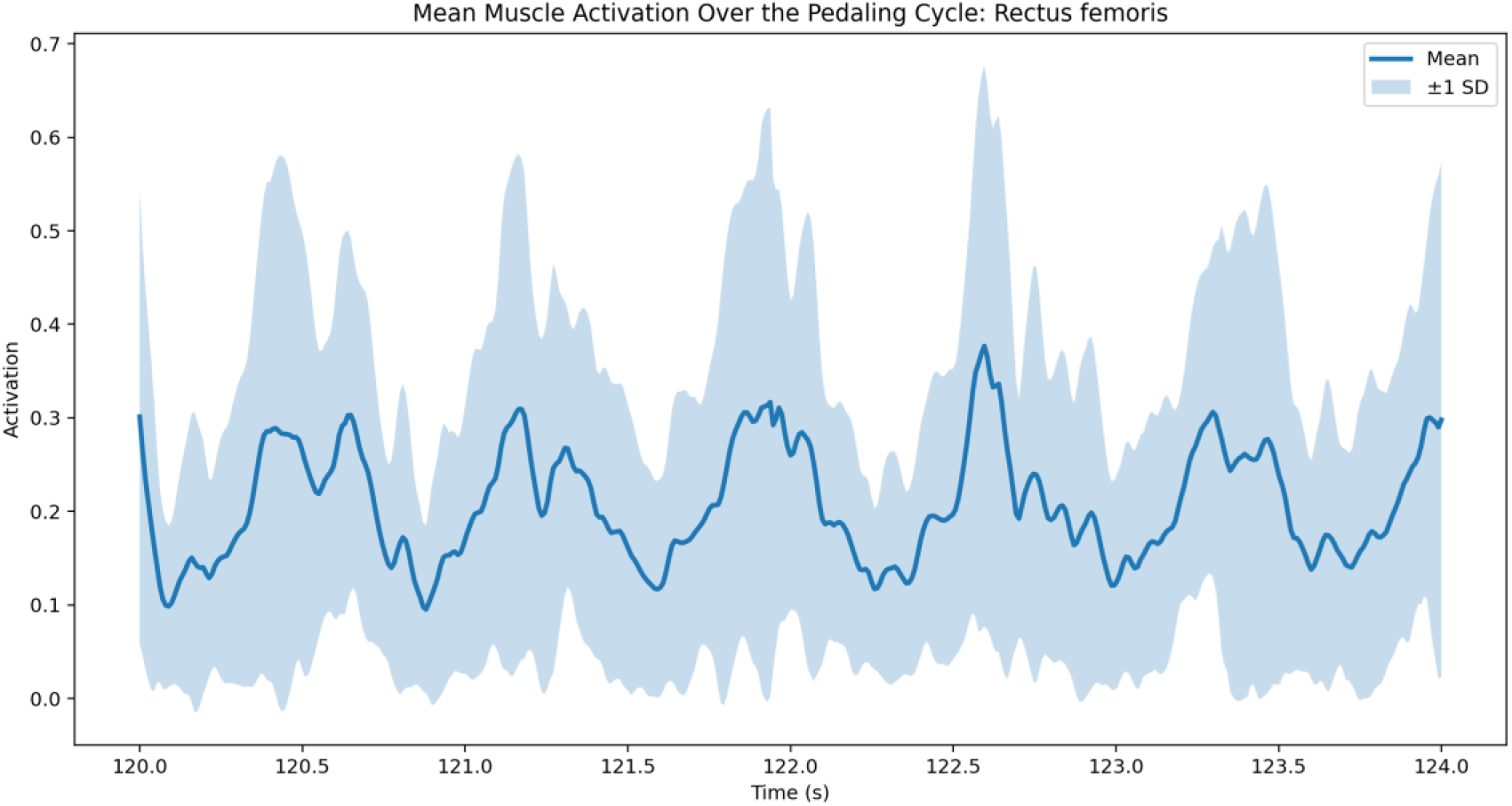
Mean ± SD activation profile of the rectus femoris.

**Appendix Figure A2.**
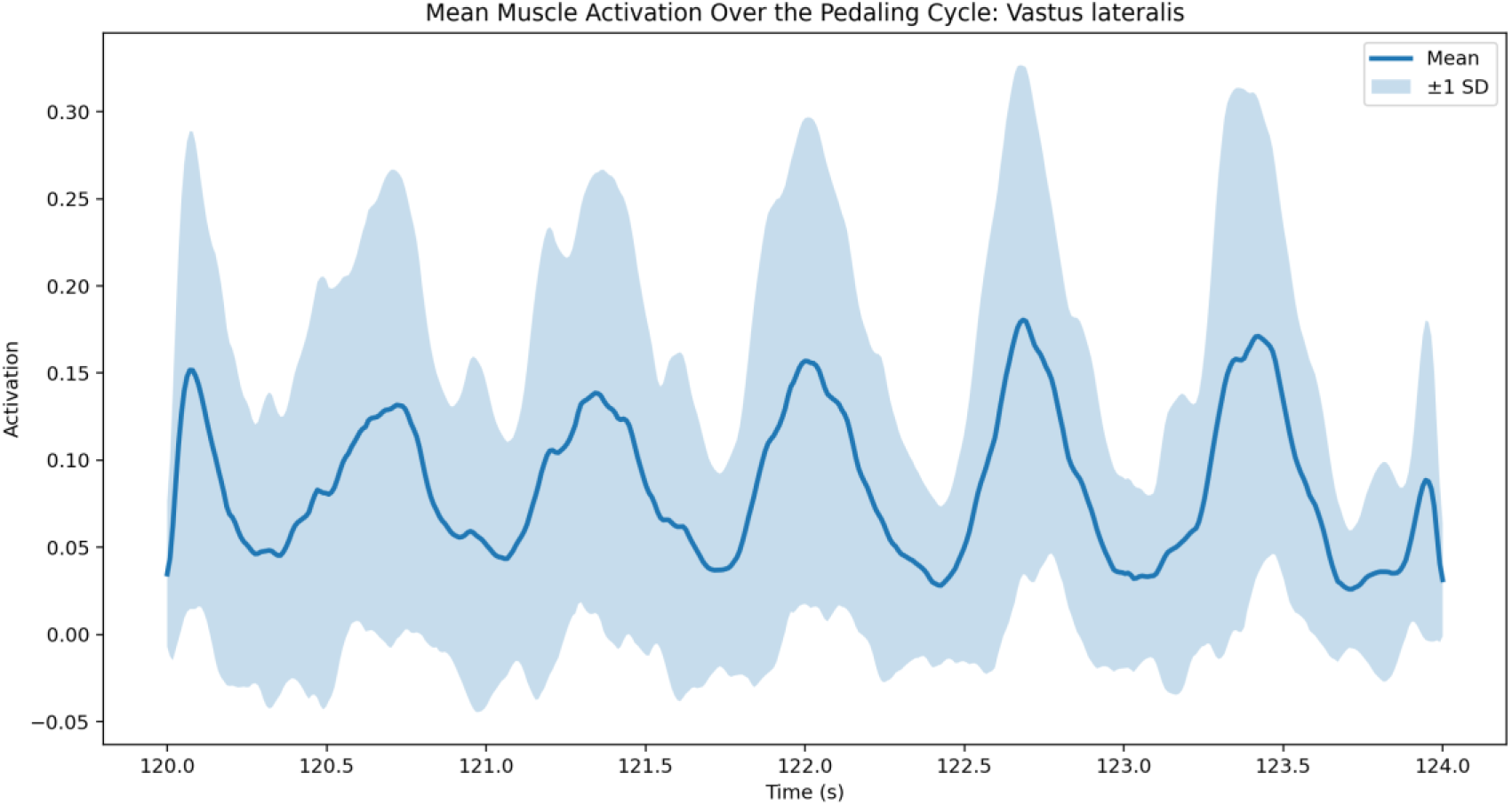
Mean ± SD activation profile of the vastus lateralis.

**Appendix Figure A3.**
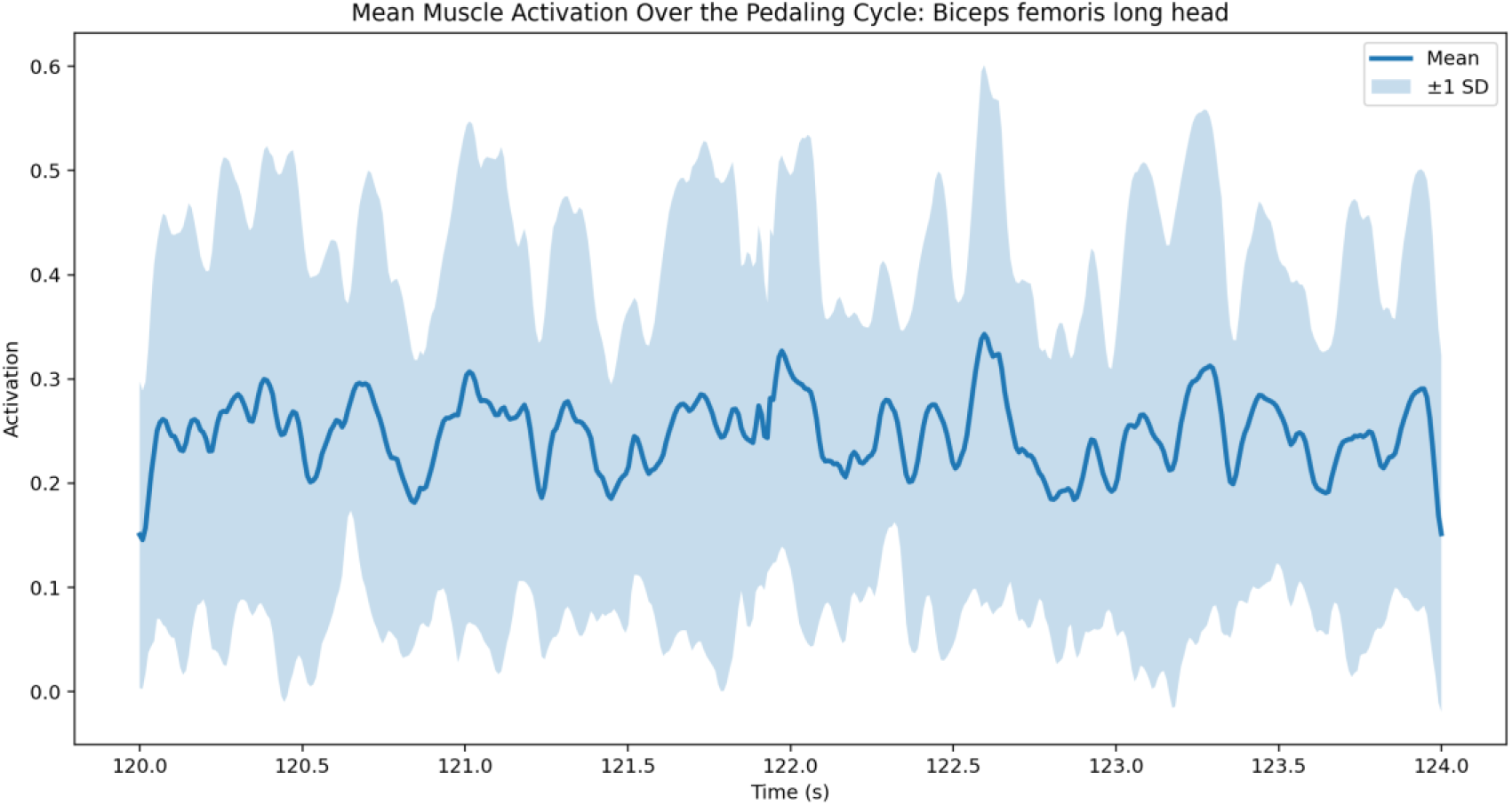
Mean ± SD activation profile of the biceps femoris long head.

**Appendix Figure A4.**
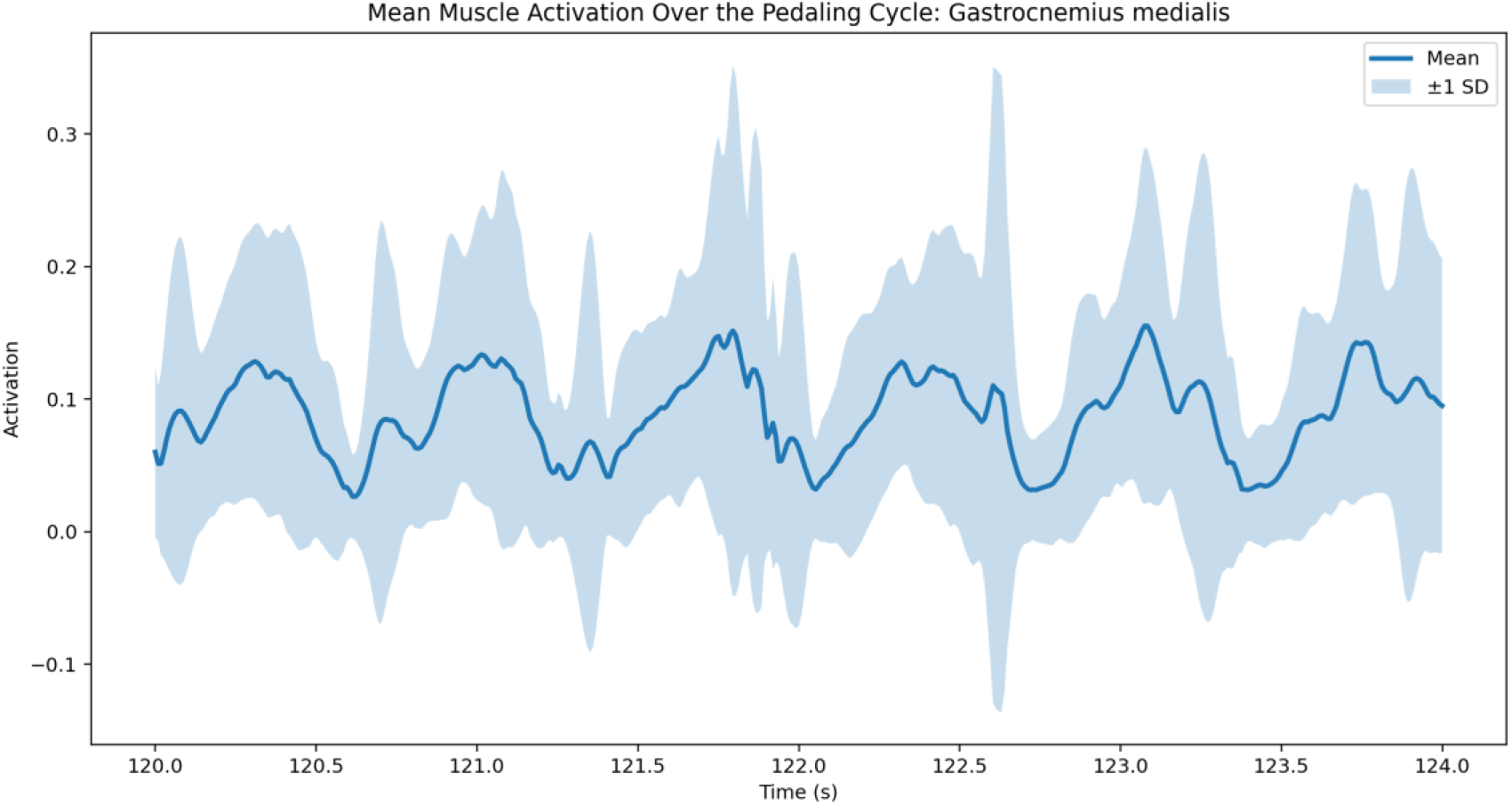
Mean ± SD activation profile of the gastrocnemius medialis.

**Appendix Figure A5.**
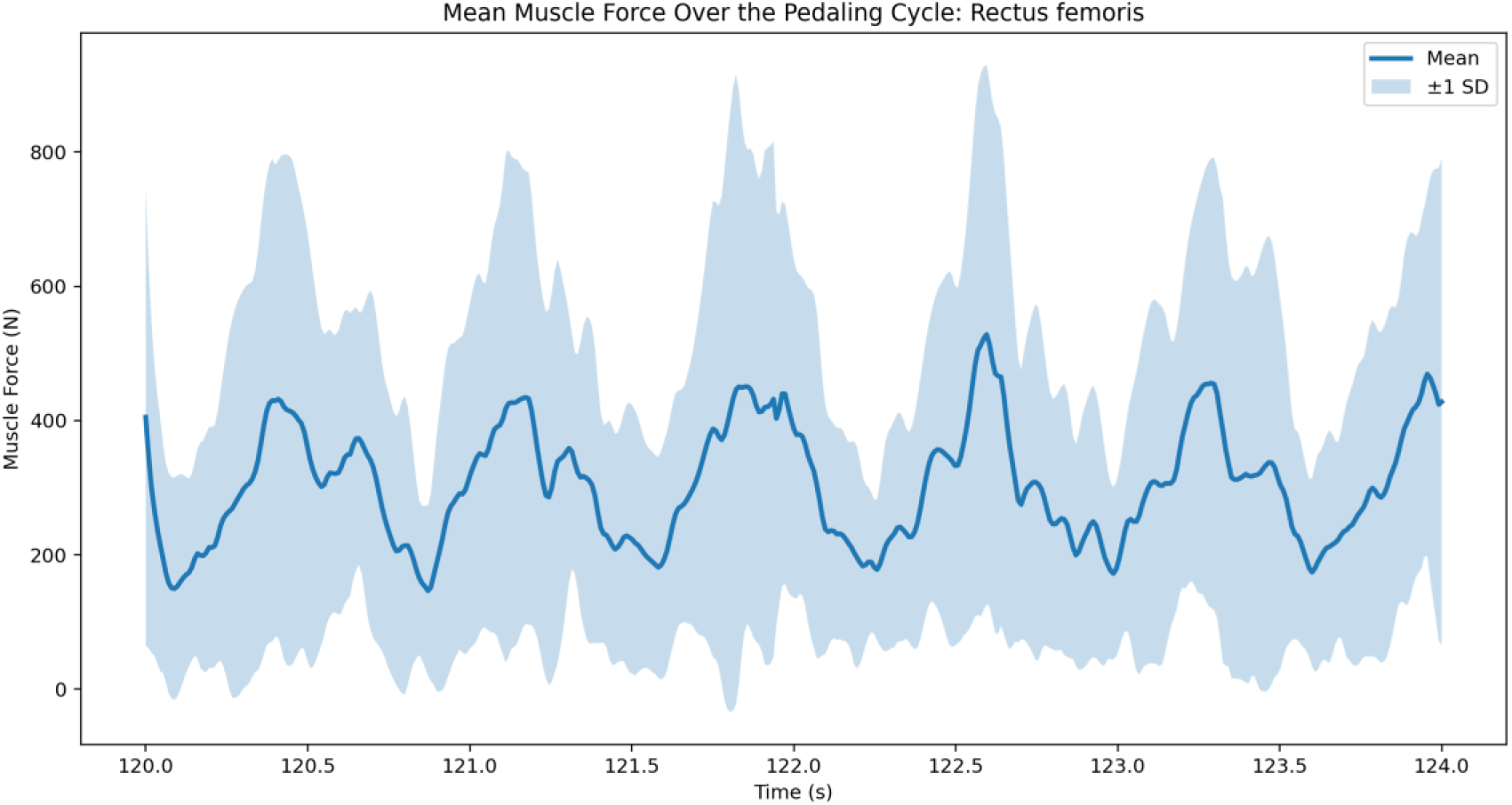
Mean ± SD force profile of the rectus femoris.

**Appendix Figure A6.**
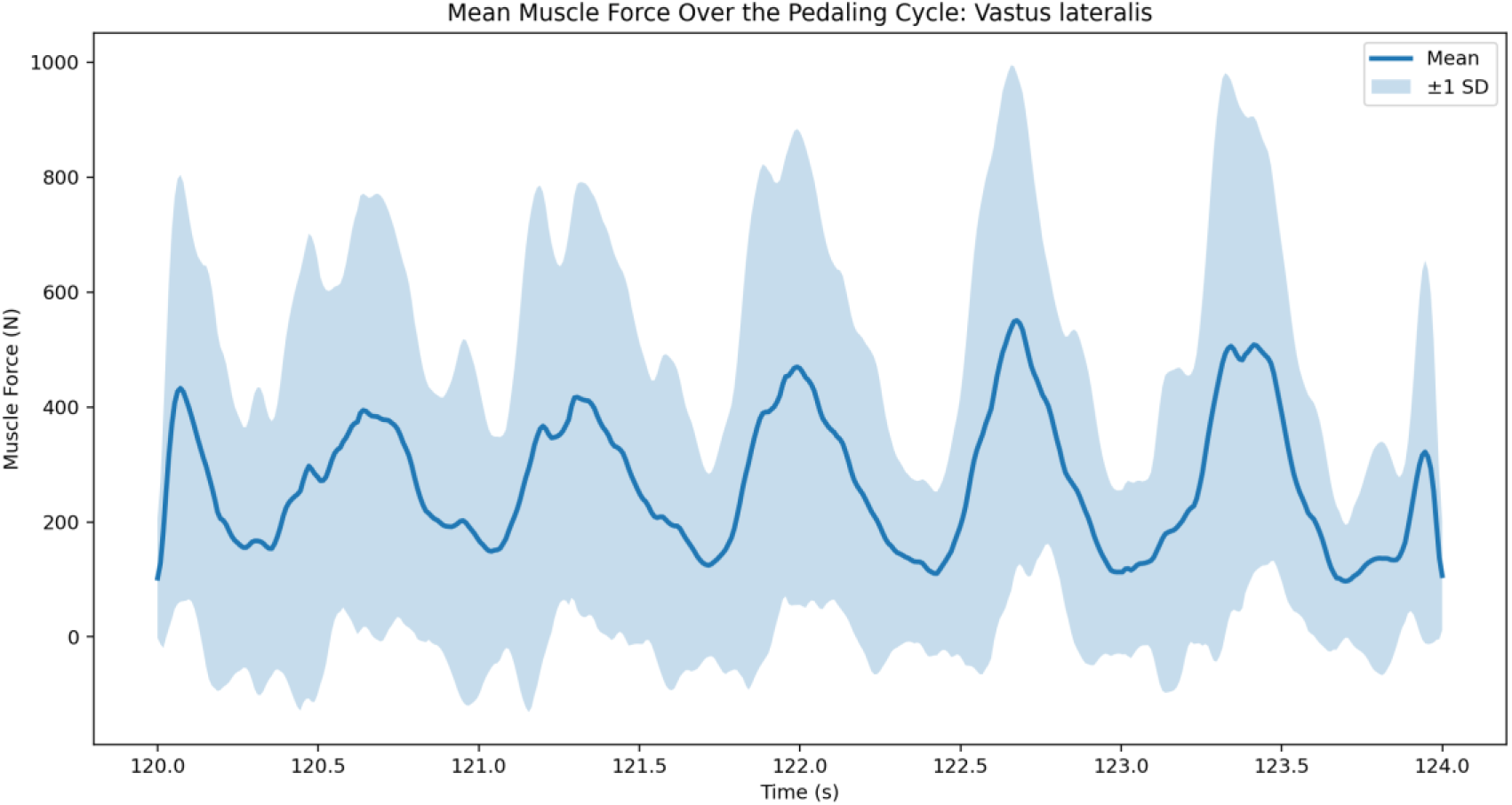
Mean ± SD force profile of the vastus lateralis.

**Appendix Figure A7.**
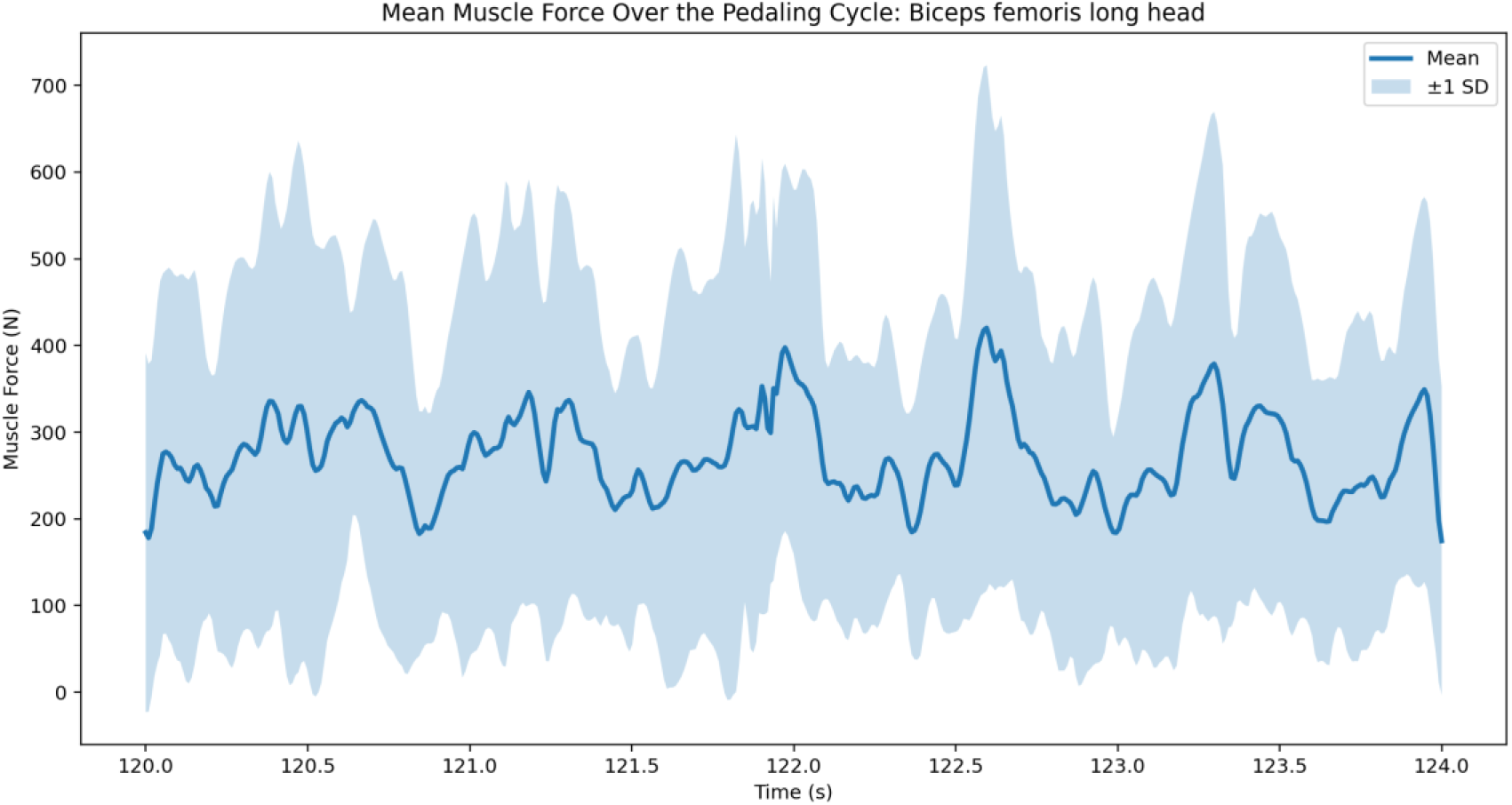
Mean ± SD force profile of the biceps femoris long head.

**Appendix Figure A8.**
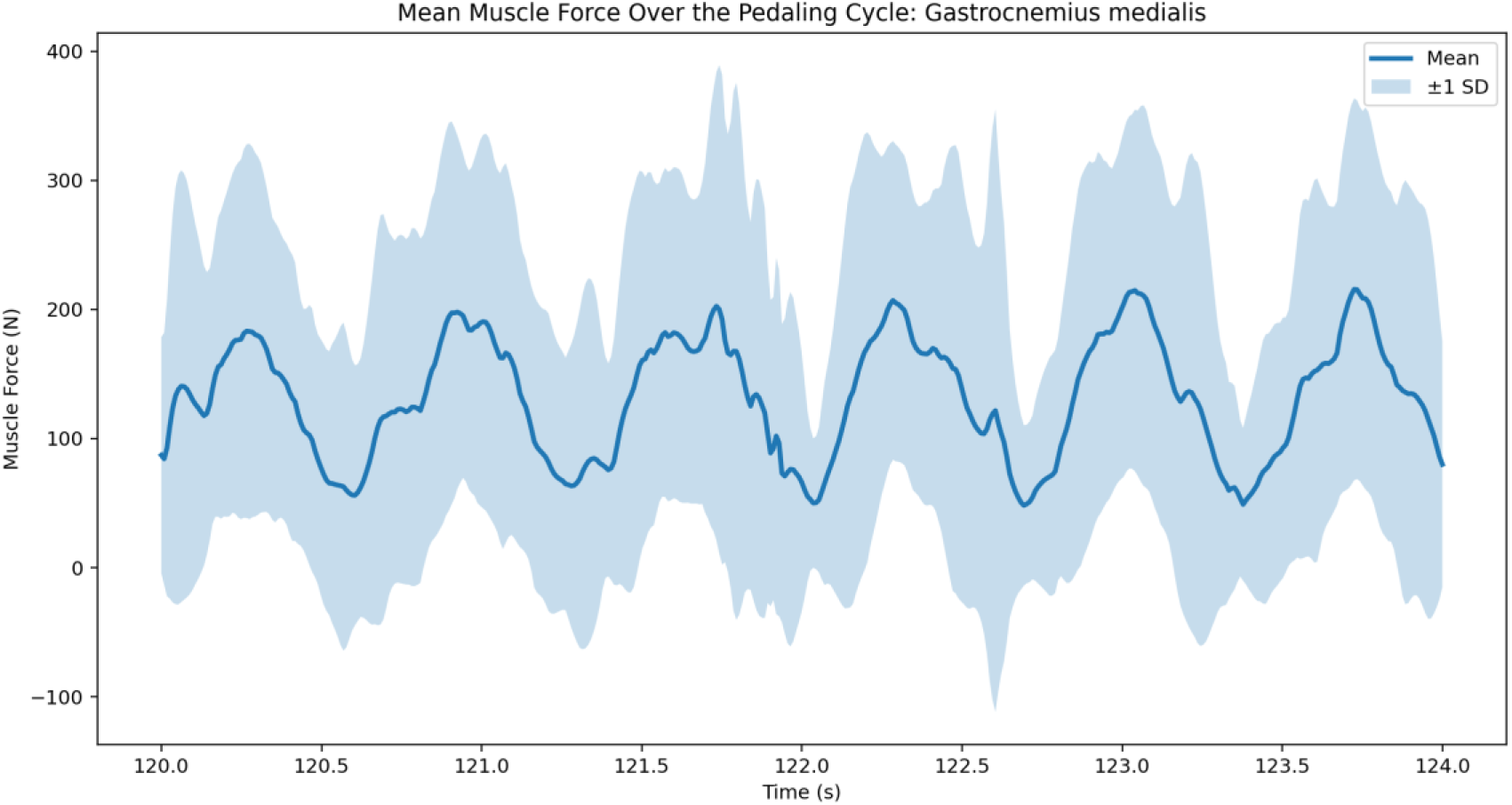
Mean ± SD force profile of the gastrocnemius medialis.

